# Bats possess the anatomical substrate for a laryngeal motor cortex

**DOI:** 10.1101/2023.06.26.546619

**Authors:** Alexander A. Nevue, Claudio V. Mello, Christine V. Portfors

## Abstract

Cortical neurons that make direct connections to motor neurons in the brainstem and spinal cord are specialized for fine motor control and learning [1, 2]. Imitative vocal learning, the basis for human speech, requires the precise control of the larynx muscles [3]. While much knowledge on vocal learning systems has been gained from studying songbirds [4], an accessible laboratory model for mammalian vocal learning is highly desirable. Evidence indicative of complex vocal repertoires and dialects suggests that bats are vocal learners [5, 6], however the circuitry that underlies vocal control and learning in bats is largely unknown. A key feature of vocal learning animals is a direct cortical projection to the brainstem motor neurons that innervate the vocal organ [7]. A recent study [8] described a direct connection from the primary motor cortex to medullary nucleus ambiguus in the Egyptian fruit bat (*Rousettus aegyptiacus)*. Here we show that a distantly related bat, Seba’s short-tailed bat (*Carollia perspicillata*) also possesses a direct projection from the primary motor cortex to nucleus ambiguus. Our results, in combination with Wirthlin et al. [8], suggest that multiple bat lineages possess the anatomical substrate for cortical control of vocal output. We propose that bats would be an informative mammalian model for vocal learning studies to better understand the genetics and circuitry involved in human vocal communication.

## Main Text

Speech learning in humans is largely based on imitation of an adult tutor [9]. This vocal learning process is thought to depend on cortical circuits and is disrupted by a number of genetic disorders ranging from FOXP2-related speech and language disorder to autism spectrum disorder [10, 11]. To fully understand the neural mechanisms of vocal learning and the underlying causes of related disorders, a mammalian vocal learning model is necessary. For decades, songbird species like zebra finches have been the laboratory model of choice to study vocal learning because they undergo a neurodevelopmental song learning process with several parallels to human speech acquisition [4]. While much has been learned about vocal learning behavior and related circuitry in songbirds, its translation to humans is not without limitations. Humans and songbirds lineages split over 300 million years ago, and have evolved different cortical organization [12]. A mammalian system for studying vocal learning may more closely parallel human speech acquisition and related circuits, and serve as means for which communication disorders can be modeled.

To date, there is limited evidence for vocal learning in non-human mammals. A number of studies have examined whether mice are vocal learners, as they are highly amenable to laboratory experiments and genetic manipulation [13-15]. While mice may be valuable models for studying vocal communication [16], they do not learn their social vocalizations [17-20]. In contrast, bats exhibit multiple behavioral hallmarks of vocal learning [6], including disruption in vocalizations from acoustic isolation [5], evidence of dialects [21], and babbling [22]. The babbling behavior in bats exhibits many features seen in humans, where it is a key component of language development [23]. Despite the behavioral studies suggesting vocal learning, little is known about the neural circuits that would enable this trait in bats. To further validate the use of bats as a model for vocal learning, it is necessary to determine whether bats have the neural circuitry required for this complex behavior.

Control of vocal output via direct motor cortex projections to brainstem motor neurons is critical to vocal learning in both humans and songbirds. In humans, the laryngeal motor cortex (LMC) projects to nucleus ambiguus [7], which contains laryngeal motor neurons, (Figure 1A) and in songbirds, the robust arcopallial (RA) nucleus projects to vocal motor neurons in a subdivision of the hypoglossal nucleus [24]. A recent study revealed that a region of the primary motor cortex in the Egyptian fruit bat (*Rousettus aegyptiacus*; Yinpterochiroptera, Pteropodidae) makes a monosynaptic projection to the vicinity of cricothyroid-projecting motor neurons [8] and is active when the bats are vocalizing. While this represents the first report of a laryngeal representation within a bat motor cortex, bats are highly speciated, with distantly related families showing behavioral evidence of vocal learning.

**Figure 1:**
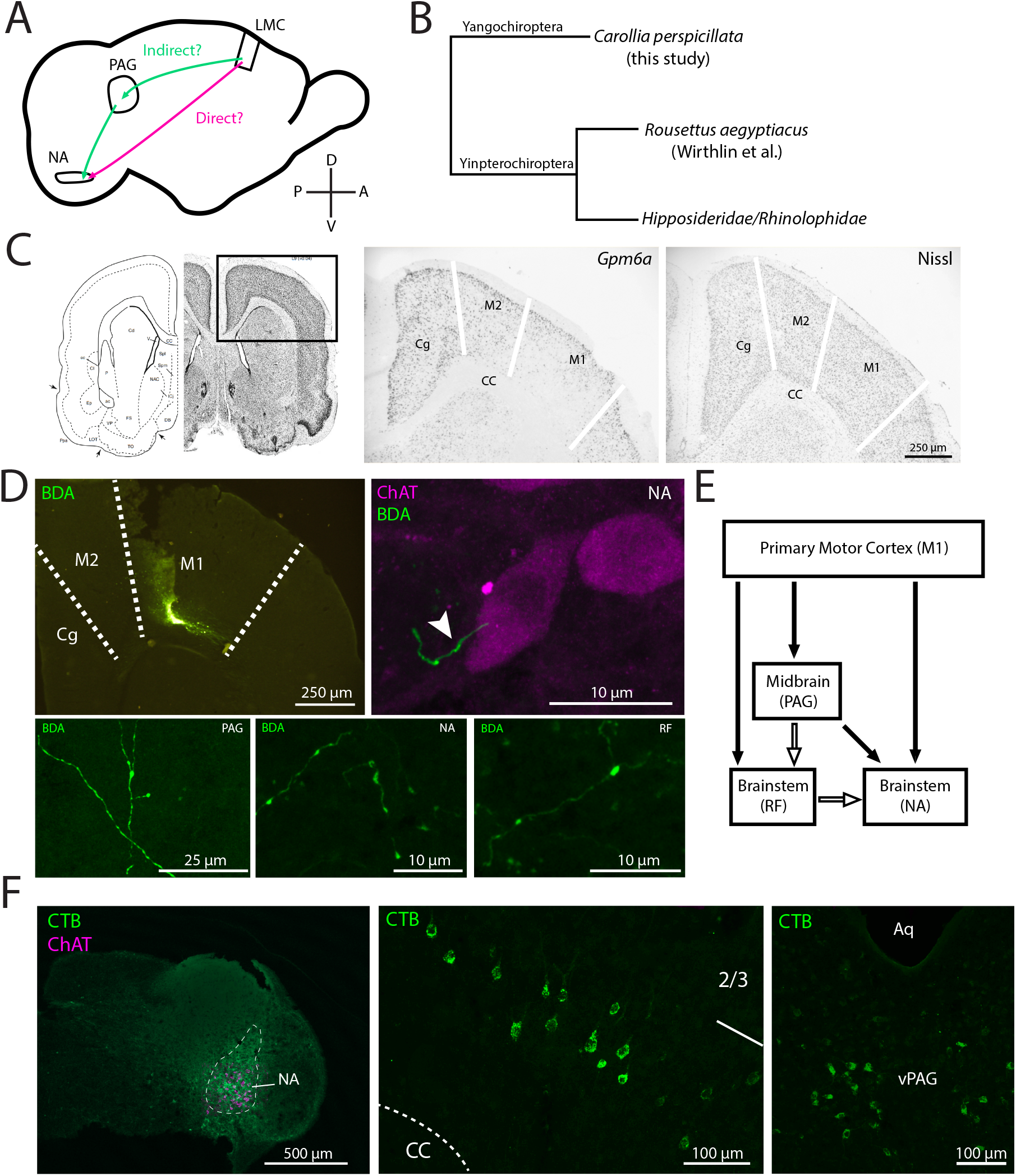
A. The direct (magenta) and indirect (green) cortical pathways for vocal control. B. A simplified cladogram of bats highlighting the laryngeal echolocating Yangochiroptera (most microbats) and the distantly related Yinpterochiroptera, which contains both non-laryngeal echolocating bats (megabats) and some microbat families. C. In situ hybridization for Gpm6a (middle) delineates the primary motor cortex (M1) in C. perspicillata. Nissl stain (right) and atlas location (left) [3]. Boundary between M2 and Cg was estimated from layer 6 GPM6A+ cells. Boundary between M2 and M1 was estimated from absence of GPM6A labeling in layer 5. D. Small, deep layer localized anterograde tracer injection into M1 (top left panel) reveals fibers with putative puncta in nucleus ambiguus, reticular formation, and periaqueductal gray (bottom panels). Fibers in nucleus ambiguus exhibit presumed synaptic boutons in close proximity to ChAT+ motor neurons (top right). E. Projections described in the present study (closed arrows) and canonical projections not assessed (open arrows). F. Retrograde tracing from a brainstem injection targeted at nucleus ambiguus (left) shows labeled cells in deep layer projection neurons in M1 (center) and in ventral periaqueductal gray (right). Abbreviations: M1: non-laryngeal motor cortex primary motor cortex; Cg: cingulate cortex; M2: secondary motor cortex; CC: corpus callosum; NA: nucleus ambiguus; RF: reticular formation; Aq: aqueduct; vPAG: ventral periaqueductal gray.

In the present study we explored whether Seba’s short-tailed bat (*Carollia perspicillata*; Yangochiroptera, Phyllostomidae, Figure 1B) possesses the anatomical substrate for a laryngeal motor cortex. *C. perspicillata* are gregarious bats with a complex social repertoire [25]. While *C. perspicillata* itself has not been studied for vocal learning capabilities, multiple species within Phyllostomidae exhibit this trait [26, 27]. In contrast to *R. aegyptiacus*, which echolocate using tongue clicks, *C. perspicillata* produce echolocation calls laryngeally [28], suggesting they may have precise control over laryngeal muscles. These behavioral characteristics support *C. perspicillata* as an informative mammalian model for understanding the neural mechanisms underlying vocal learning and associated communication disorders.

To search for evidence of an LMC in *C. perspicillata*, we first utilized *in situ* hybridization to identify the primary motor cortex based on expression of *Gpm6a*. While this gene was reported to share convergent downregulation in human LMC and songbird RA [29], we have previously shown that *Gpm6a* downregulation in songbirds is not limited to RA but also includes the adjacent arcopallial motor region, analogous to the non-laryngeal representation of the primary motor cortex in mammals [30]. The boundaries of primary motor cortex (M1) in *C. perspicillata* could be delineated with the *Gpm6a* expression pattern, noting the downregulation in deep layers (Figure 1C).

We then investigated whether *C. perspicillata* possess a direct projection from motor cortical neurons to nucleus ambiguus, a characteristic feature of vocal learners [7]. We injected an anterograde tracer targeted to the deep layers of the primary motor cortex area identified by gene expression (Figure 1D, upper left panel), and found labeled fibers in nucleus ambiguus, as well as in the periaqueductal grey (PAG) and reticular formation (Figure 1D, bottom panels). Upon staining for choline acetyltransferase (ChAT) to label brainstem motor neurons, we found that the anterogradely labeled fibers from the motor cortical injection were in close proximity of ChAT+ cells in nucleus ambiguus (Figure 1D, upper right panel). To confirm this projection, we injected a retrograde tracer into the nucleus ambiguus (Figure 1F, left panel), which resulted in labeled cell bodies both in the primary motor cortex overlapping the *Gpm6a* downregulation zone and in the ventrolateral PAG (Figure 1F, middle and right panels), noting the tracer was not restricted to nucleus ambiguus. These results provide supportive evidence of a direct projection from the primary motor cortex to vocal motor neurons in nucleus ambiguus in *C. perspicillata* (Figure 1E). While the fibers terminating in nucleus ambiguus represent a direct cortical projection that may support fine motor control of laryngeal muscles, the apparent projections from motor cortex to PAG and reticular formation, and from these areas to nucleus ambiguus suggest that *C. perspicillata* may also possess an indirect cortical vocal pathway, which could possibly be involved in non-learned vocalizations [31]. Overall, these results support the conclusion that *C. perspicillata* possess the neural circuitry necessary for precise control of vocalizations.

A direct projection from the motor cortex to the nucleus ambiguus is not present in non-vocal learning animals including non-human primates, cats, rats, and non-vocal learning birds [7], though there is a reported sparse projection from motor cortex to nucleus ambiguus in mouse [32]. However, mice vocalizations are innate, not learned [17, 18], and mice produce ultrasonic vocalizations (USV) through a different mechanism than humans produce speech [33], suggesting that mice USVs may not be under the same degree of cortical control. Recent assessments have proposed that vocal learning is not binary, where a species either can or cannot learn via imitation, but rather that vocal learning is on a spectrum [34] and may involve several modules that could reflect partial components of vocal learning systems [35]. It is unclear as of yet where bats fall along this spectrum of vocal learners, how many bat species are capable of vocal learning, or whether different species of bats exhibit a greater degree of vocal learning than others [36].

Bats likely produce echolocation calls and vocalizations using different laryngeal muscles [37], but duplication and specialization of the echolocation pathway may serve as a possible origin for vocal learning abilities. In combination with Wirthlin et al. [8], our findings point to Pteropodidae and Phyllostomidae as the first non-human mammalian groups shown to exhibit both behavioral hallmarks and brain anatomy for vocal learning. While this suggests that the anatomical projection was present in the common ancestor of all bats, with the less parsimonious explanation being multiple independent gains, characterizing more bat species is needed to further clarify the origin of bat vocal learning.

In summary, this study adds to the evidence that bats possess the anatomical substrate to produce learned vocalizations. Bats are the only non-human mammal known to exhibit both the behavior and neural circuitry for vocal learning. Our results, in combination with the sociality of bats, suggest that bats are an accessible model to study mammalian imitative vocal learning, and subsequently diseases and disorders that affect vocal communication in humans.

## Methods

### Animal and tissue preparation

We used adult Seba’s short-tailed bats (*Carollia perspicillata*) bred in our colony. All animals lived in a flight room under natural harem social structures and had free access to food and water. All care and procedures were in accordance with the guidelines of the National Institutes of Health and were approved by the Washington State University Institutional Animal Care and Use Committee.

### In situ hybridization

Prior to tissue collection, bats (n=4, males) were isolated overnight in a custom-built sound-proof chamber to reduce auditory stimulation. A microphone was placed near the cage to ensure the bats were not vocalizing for 1-2 hours prior to tissue collection. Bats were sacrificed after isoflurane inhalation and brains were rapidly removed, blocked coronally, and frozen with OCT (Tissue-Tek) in a dry ice/isopropyl alcohol slurry. Brains were sectioned coronally at 10μm with a cryostat (Leica) and stored at -80°C until use. Nissl staining was performed on adjacent sections to those used for *in situ* hybridization. We used a cDNA clone from the Mammalian Gene Collection for GPM6A (Human, Clone ID #BI670057). The protocol for *in situ* hybridization has been described previously [38]. Briefly, the clone was digested and purified to recover cDNA insert. Digoxygenin- (DIG) labeled antisense riboprobes were generated from the cDNA template using T7 RNA polymerase and a DIG-UTP RNA labeling kit (Roche). Brain sections were fixed in 3% phosphate buffered paraformaldehyde for 5 min followed by a rinse in phosphate buffered saline. Sections were then acetylated for 10 min (0.25% acetic anhydride in 1.4% triethanolamine and dH2O), washed in 2X SSPE, and dehydrated through an ethanol series. Sections were incubated in hybridization solution (50% formamide, 2X SSPE, 2 μg/μL tRNA, 1μg/μL BSA,1μg/μL Poly A, and 2 μL of riboprobe per slide), coverslipped and incubated overnight in mineral oil at 62°C. The following day, slides were washed in chloroform and de-coverslipped in 2X SSPE. Slides were then washed in post-hybridization washes at 62 °C, first in 50% formamide in 2X SSPE for 70 min, then two washes in 0.1X SSPE for 30 min with agitation every 10 min. Slides were then transferred to a solution of 0.3% Triton X-100 in a Tris-HCl, NaCl buffer, sections were framed with a PAP pen, and blocked with 8.3 ng/μL of BSA in a Tris-HCl, NaCl buffer (TNB) for 30 min. Slides were then incubated in alkaline phosphatase-labeled anti-DIG antibody (Roche; 1:600) in TNB for 2 h. Slides were washed in Tris-HCl, NaCl, and MgCl2 buffer, placed in slide jars, and incubated in 5-bromo-4-chloro-3-indolyl-phosphate/nitroblue tetrazolium (BCIP/NBT; Perkin-Elmer) at room temperature overnight. Following chromogen incubation, slides were washed in dH2O and fixed in 3% phosphate-buffered paraformaldehyde for 5 min. Slides were air-dried and coverslipped with VectaMount AQ (Vector Labs).

### Tract tracing and immunohistochemistry

For tracer injections, bats were anesthetized by isoflurane inhalation in a custom stereotaxic apparatus. A 1mm x 1mm craniotomy was made over the primary motor cortex. A tracer deposit via iontophoretic injection (5μA, 10 min) at varying depths was made targeting the deep layers of the primary motor cortex using 10% biotinylated dextran amine (BDA; 10,000 MW) (Life Technologies) (n=4, males) or 1% cholera toxin subunit B (CTB, List Biological Laboratories) in nucleus ambiguus/brainstem (n=2, males). We then covered the craniotomy with petroleum jelly and bone wax to prevent the brain from dehydrating, applied lidocaine and Neosporin to the exposed tissue, and returned the bat to a home cage. After one week the bat was deeply anesthetized with isoflurane and transcardially perfused with 10% buffered formalin. The brains were dissected and cryoprotected overnight in 20% sucrose in 0.1M phosphate buffer (PB). We sectioned coronally at 40 μm using a Leica freezing microtome.

The primary antibody solution contained of goat anti-ChAT (Millipore AB144P, 1:200) and anti-CTB (List Biological Laboratories, 1:10,000) antibodies, 3% normal donkey serum (Millipore), and 0.4% Triton X-100 (Sigma-Aldrich) in 0.1M PB. Primary antibodies were visualized using Alexa Fluor 488- and 568-conjugated secondary antibodies (1:500; Life Technologies) and BDA was visualized using Alexa Fluor 568-conjugated streptavidin (Life Technologies; 1:250). Sections were then mounted on slides, dehydrated and cleared, and then coverslipped with DPX (Electron Microscopy Sciences). Label was observed using a Leica TCS SP8 confocal microscope. Brightness and contrast adjustments were used to enhance images.

## Acknowledgments

We would like to thank Peter Lovell and Alexandra Wilmington for help with *in situ* hybridization experiments. This project was supported by NIH R21HD094410 to CVP and NIH GM120464 and NSF IOS1456302 to CVM.

## Author Contributions

AAN, CVM, and CVP conceived the project and designed experiments. AAN performed experiments. AAN wrote the paper with input from CVM and CVP.

